# Optimized protocol for processing murine tumor-bearing lung tissue for flow cytometry and single cell RNA-sequencing

**DOI:** 10.1101/2025.11.25.690629

**Authors:** Sicong Wang, Anastasiia Ivanova, Helen P. Makarenkova, Katja A. Lamia

**Affiliations:** The Scripps Research Institute, Department of Molecular and Cellular Biology

## Abstract

NSCLC accounts for approximately 85% of lung cancers, and Kirsten rat sarcoma (KRAS) is the most commonly mutated gene in NSCLC. Although management of NSCLC has advanced markedly in recent years, its heterogeneity and plasticity remain incompletely understood. Single-cell RNA sequencing (scRNA-seq) has become an essential tool for dissecting cellular heterogeneity in complex tissues such as the lung. However, generating high-quality single-cell suspensions from tumor-bearing lung tissue is challenging, particularly during tumor formation when the tissue becomes fibrotic and heterogeneous. A key requirement for successful scRNA-seq is the preparation of viable single cells with minimal processing-induced stress or damage. Here, we present a detailed step-by-step protocol for isolating high-quality single-cell suspensions from both healthy and tumor-bearing mouse lung tissue in a Kras^G12D^; tdTomato reporter mouse model. The procedure includes tissue perfusion, controlled enzymatic digestion, gentle mechanical dissociation, red blood cell lysis, filtration, and viability assessment, and achieves >85% viable cells prior to scRNA-seq. Finally, the tdTomato^+^ tumor cells are efficiently isolated by FACS and can be detected as distinct clusters by scRNA-seq using this protocol.

**SUMMARY:** We present a detailed protocol for isolating high-quality single-cell suspensions from both healthy and tumor-bearing murine lung tissue. This protocol can be applied to scRNA-seq, flow cytometry analysis, and primary cell isolation.

## INTRODUCTION

Lung cancer is one of the leading causes of cancer-related deaths worldwide. It can be broadly classified into two major subtypes: small-cell lung cancer (SCLC) and non-small-cell lung cancer (NSCLC)^1^. NSCLC accounts for approximately 85% of all lung cancer cases, with adenocarcinoma being the most common histological subtype^2^. Although therapeutic strategies for NSCLC have improved substantially over the past decades, the mortality rate of lung cancer remains high^3^. Chemotherapy and immunotherapy are the two main treatment modalities used in clinical practice; however, many patients experience drug resistance or fail to respond to immunotherapy^4,5^.

The tumor microenvironment (TME) plays a pivotal role in tumor initiation, progression, and therapeutic resistance^6^. Increasing evidence suggests that the composition of the TME changes dynamically over time^7,8^. Anti-tumor immune cells are typically enriched in the TME during the early stages of tumorigenesis, whereas immunosuppressive and cancer-associated cell populations become predominant as the tumor develops. Therefore, understanding the heterogeneity and plasticity of the TME during lung cancer progression is essential for elucidating disease mechanisms and improving therapeutic outcomes.

Single-cell RNA sequencing (scRNA-seq) enables high-resolution profiling of gene expression at the individual cell level, allowing comprehensive characterization of complex tissue transcriptomes^9^. This technique has become indispensable for studying cellular heterogeneity and identifying mechanisms underlying tumor progression and drug resistance in lung cancer^10^. Such insights have facilitated the development of novel therapeutic strategies. A successful scRNA-seq experiment generally involves three key components: (1) the use of an appropriate heterogeneously multicellular experimental system, (2) the preparation of a high-quality single-cell suspension, and (3) data analysis. Among these, the process of single-cell suspension preparation is particularly critical, as cell stress or damage during dissociation can lead to inaccurate results ^11,12^.

This protocol aims to efficiently dissociate murine lung tissue while minimizing processing-induced cell stress and damage. Optional fluorescence-activated cell sorting (FACS) steps are included for quality control and enrichment of specific cell populations. Additional guidance is provided for antibody staining, DAPI labeling, and short-term fixation procedures when immediate sorting is not feasible.

## PROTOCOL

NOTE: nine or ten weeks old Kras^LSL-G12D/+^; TdTomato^+/-^ mice were used as lung cancer models. They are infected intratracheally with lentivirus expressing Cre recombinase. Tumor-bearing lung tissues were collected after 25 weeks.

### 1. Preparation of Buffers

1. DMEM/F12 medium: Prepare 2% (v/v) heat-inactivated FBS in DMEM/F12 medium.
2. Cell wash buffer: Prepare 2% (v/v) heat-inactivated FBS in Ca+/Mg+ free PBS (pH 7.4).
3. Perfusion buffer: Pre-chilled Ca^+^/Mg^+^ free PBS (pH 7.4).
4. Enzyme stock solutions: Dissolve 10 mg of collagenase IV (16000 U/mL) in 1 mL DMEM/F12 medium; Dissolve 42 mg of Dispase II (50 U/mL) in 1 mL DMEM/F12 medium; Dissolve 5 mg of DNase I (5 mg/mL) in DMEM/F12 medium.
5. Digestion medium: Mix 250 µL of collagenase IV (800 U/ml), 200 µL of Dispase II (2 U/mL), and 100 µL of Dnase I (0.1 mg/mL) in 5 mL DMEM/F12 medium.
6. Resuspension buffer: Mix 1 µL of 0.5 M EDTA (10 µM), 1 mL heat-inactivated FBS (2% v/v) in 50 mL Ca^+^/Mg^+^ free PBS (pH 7.4).

### 2. Collection of murine lungs tissue

1. Euthanize mouse in a CO_2_ chamber. Place mouse on its back on a dissection pad and pin the feet. Spray liberally with 70% ethanol.
2. Make an incision through the skin and muscles of the lower abdomen. Carefully open the thorax and avoid cutting the lungs or the heart.
3. Attach a safety-multifly 21G needle to 50 mL syringe and aspirate 30 mL perfusion buffer.
4. Insert needle in the left ventricle and cut a small incision in the right atrium using the sharp straight scissor.
5. Perfuse whole body with 15 mL perfusion buffer. NOTE: The whitening of the liver indicates that whole-body perfusion has been completed.
6. Remove the lungs from the thorax and cut away excess tissue attached to the lungs. Place the lungs in 10 mL pre-cooled Ca^+^/Mg^+^ free PBS to wash away excess blood.
7. Transfer the lungs into a 1.5 mL Eppendorf tube containing 500 µL of ice-cold DMEM/F12.

### 3. Preparation of the lung single-cell suspension

1. Cut the lung tissue into small pieces with an extra narrow straight scissor.
2. Transfer the minced lung tissue to a 6 cm dish containing 5 mL digestion medium.
3. Place 6 cm dish on a thermostatic shaker at 50 rpm for 25 min at 37℃. Mix the suspension using a 1 mL pipette every 8 min.
4. Place 100 µm mesh strainer in a 10 cm dish. Pre-wet the strainer with 1 mL of DMEM/F12 medium.
5. Transfer the digested suspension into the strainer. Push the tissue through the strainer with the rubber end of a sterile syringe plunger.
6. Place a 70 µm mesh strainer over a 50 mL centrifuge tube. Pre-wet the strainer with 1 mL of DMEM/F12 medium.
7. Filter the cell suspension through strainer and rinse the strainer with 5 mL DMEM/F12 medium.
8. Centrifuge filtered cell suspension at 400 × *g* for 10 min and remove the supernatant carefully. Resuspend the cell pellet in 1 mL Red Blood Cell Lysing Buffer for 2 min at room temperature.
9. Add 14 mL cell wash buffer and centrifuge at 400 ×*g* for 10 min. NOTE: If the pellet is still red after washing, repeat step 8 and 9.
10. Discard the supernatant and resuspend the cells in 20 mL resuspension buffer.
11. Mix 10 µL of single-cell suspension with equal volume of trypan blue and add 10 µL of mixture to cell counting chamber slide. Check live cell viability, cell number, and single-cell status using cell counter.

### 4. Flow cytometry analysis

1. Resuspend cells in ice-cold FACS buffer and transfer 100 µL of cell suspension into 1.5 mL Eppendorf tubes containing 1 µL Fc block antibody. Incubate samples for 15 min on ice.
2. Centrifuge samples at 500 × *g* for 5 min and discard the supernatant. Resuspend cells in antibody mixture and incubate 45 min in dark at 4℃.
3. Centrifuge samples at 500 × *g* for 5 min and discard the supernatant. Resuspend cells in FACS buffer containing DAPI stain (1:5000).

## REPRESENTATIVE RESULTS

High cell viability was achieved in the single-cell suspension prepared using this optimized protocol. Changes that improved cell viability for tumor-bearing lungs included reducing the concentration of enzymes and shaking and mixing the samples during incubation at 37 °C. Furthermore, we eliminated the use of an additional 40 µm mesh strainer in step 6 to reduce mechanical shear stress that increases the likelihood of cell damage; including the additional strainer may reduce multiplet formation. On average, cell viability exceeded 85%, compared to less than 80% before optimization (Table 1). Variations in cell concentration were primarily attributed to differences in lung size and the suspension medium used across replicates.

Two whole-lung single-cell suspensions were subjected to scRNA-seq on the 10x Genomics platform. Cell Ranger quality assessment identified 23,488 and 21,436 estimated cells in the two samples, respectively (Table 2). The barcode plots indicated minimal ambient RNA contamination, likely reflecting the high cell viability (Figure 1A and B).

**Figure 1.**
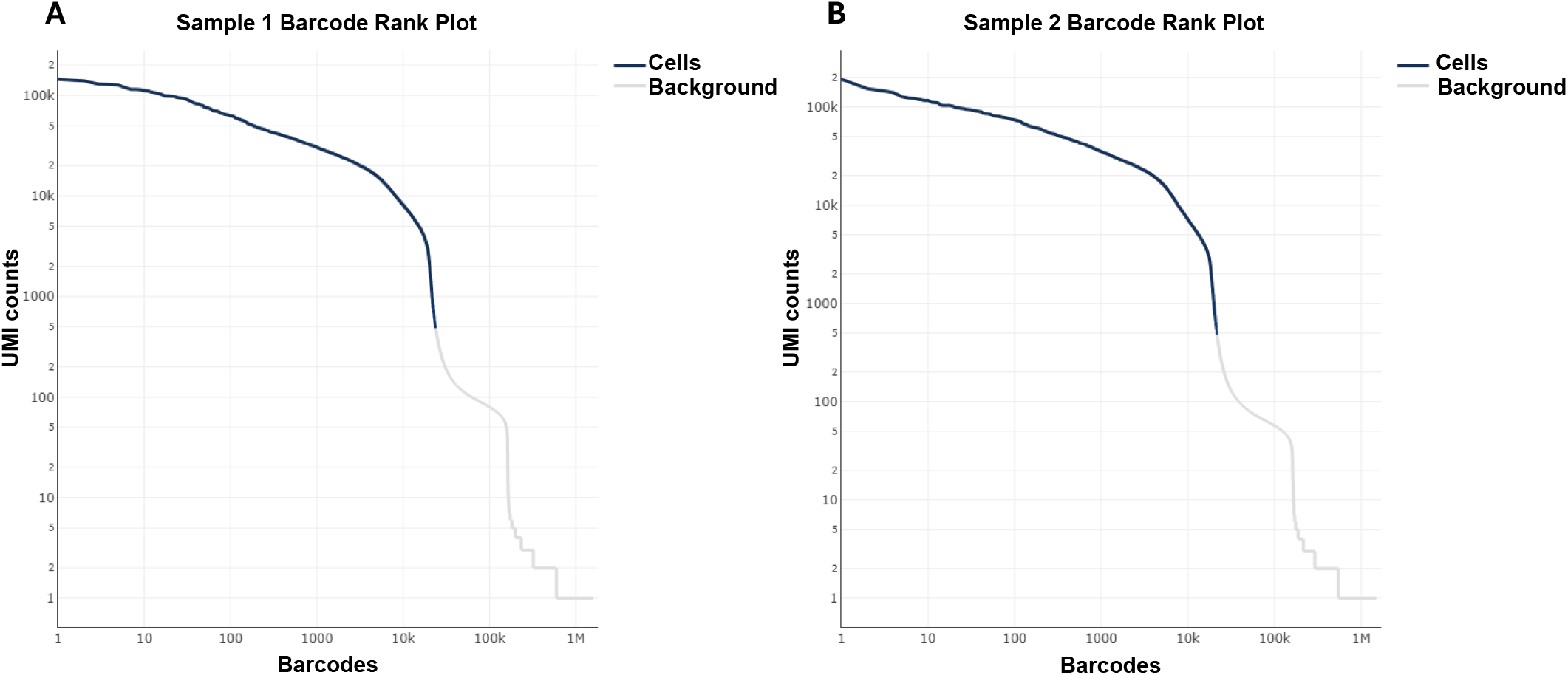
ScRNA-seq Barcode rank plot of murine tumor-bearing lungs. Barcode rank plot of two murine tumor-bearing lungs samples.

Principal component analysis (PCA) and uniform manifold approximation and projection (UMAP) revealed well-defined clusters, with all major lung cell types represented (Figure 2A). Dot plots of representative marker genes confirmed that canonical biomarkers were highly expressed in their corresponding cell types (Figure 2B). Furthermore, TdTomato^+^ cells were enriched in epithelial-related clusters, and TdTomato^+^/EpCAM^+^ cells were successfully isolated by FACS (Figure 3).

**Figure 2.**
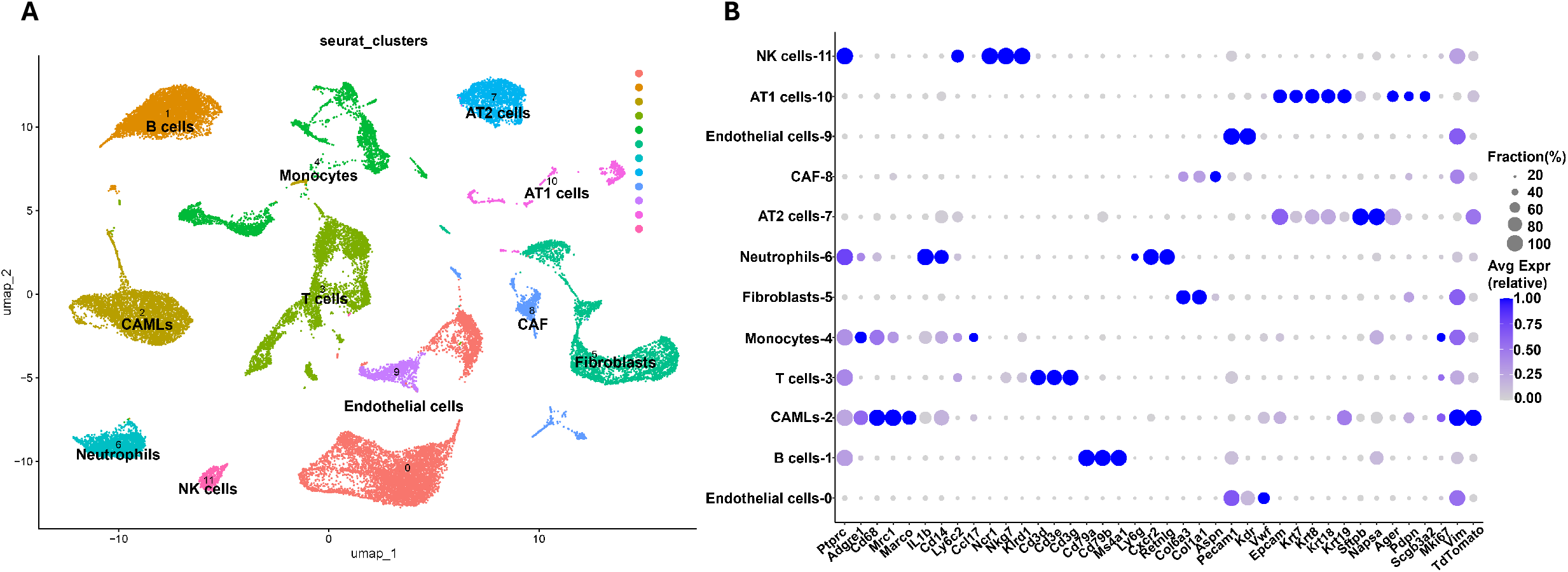
Single-cell transcriptomics analysis of murine tumor-bearing lungs. **A**. UMAP projection of whole tumor-bearing lungs datasets. **B**. Dot plot of classical genes used for cluster annotations.

**Figure 3.**
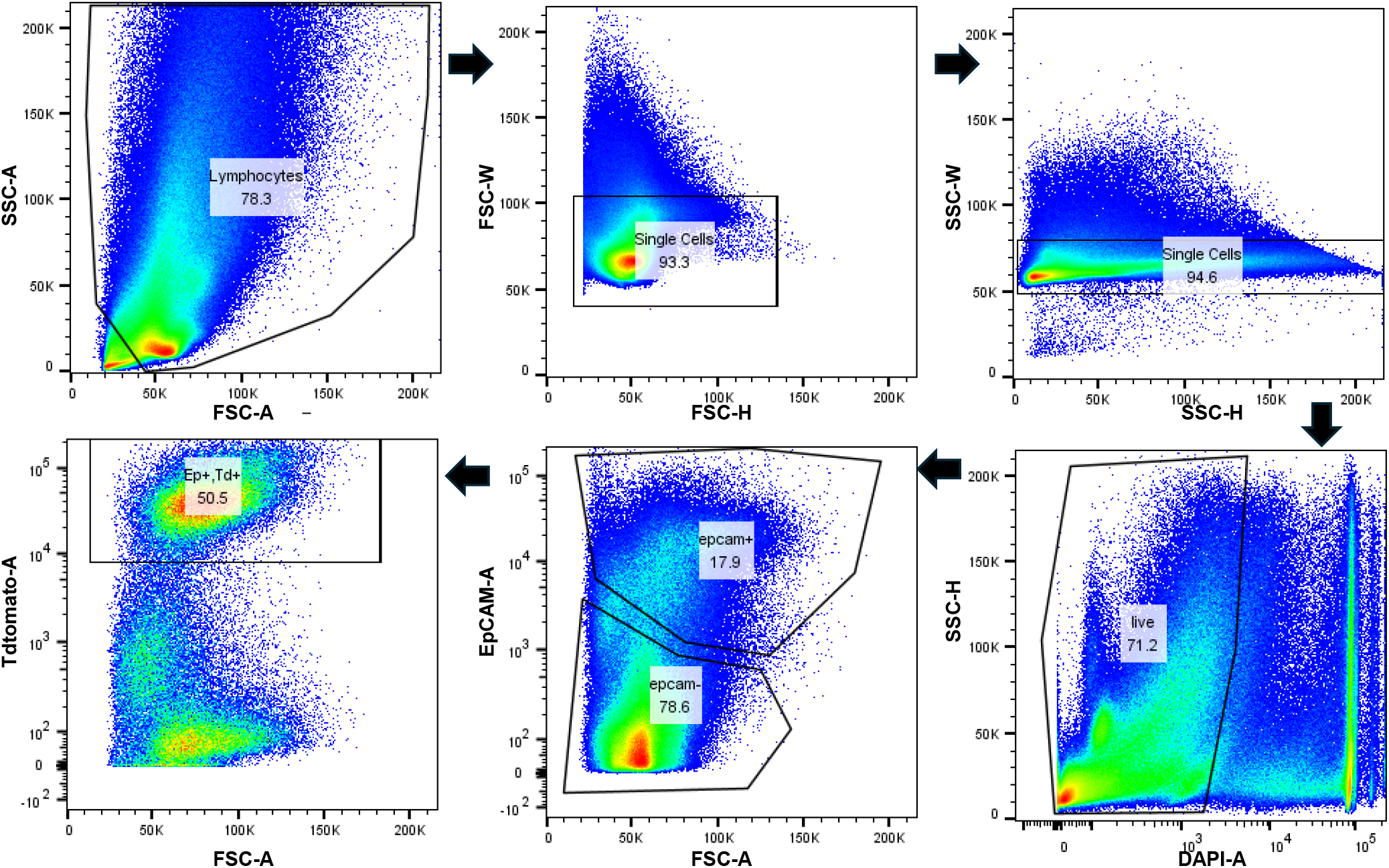
Isolating TdTomato^+^/EpCAM^+^ cells by flow cytometry. Flow cytometry gating strategy for murine lung single-cell suspension.

In summary, this optimized protocol enables efficient dissociation of murine tumor-bearing lung tissue, yielding high cell viability, low RNA contamination, and comprehensive cellular representation.

## FIGURE AND TABLE LEGENDS

**Table 1. Tumor-bearing lung single-cell suspension cell viability before and after optimizing.** The cell concentration and percentage of live cells determined before and after using this protocol.

**Table 2. ScRNA-seq quality control analysis by Cell Ranger.**x The key estimates of the single-cell data obtained from Cell Ranger

## DISCUSSION

This protocol provides a detailed step-by-step procedure for preparing high quality single cell suspensions from murine lung tissue. Several critical steps are required to achieve optimal results.

Prior to enzymatic digestion, whole body perfusion must be performed to remove blood from the lungs. This ensures that immune cells detected by scRNA-seq originate from lung tissue rather than circulating blood. During perfusion, it is essential to insert the needle into the left ventricle to prevent the perfusate from entering the pulmonary circulation and damaging lung cells. To preserve cell viability, all buffers including wash, resuspension, and perfusion solutions should be prechilled on ice. Given the inherent heterogeneity of lung tissue^13^, samples should be maintained in DMEM/F12 medium before and during digestion, as this medium provides essential nutrients that sustain cell viability.

The selected type and concentration of the collagenase/dispase mixture have been shown in multiple studies to be optimal for lung tissue digestion^10,14,15^. High cell viability and well-defined cell clusters observed in this protocol confirm that the enzyme combination is both mild and effective. Calcium and magnesium ions are known to promote cell clumping and enhance enzymatic activity^16,17^. Therefore, Ca^+^/Mg^+^ free PBS is used to prepare most buffers to minimize clumping, while DMEM/F12 containing these ions is used as the basal medium for the digestion buffer to maintain balanced enzymatic efficiency.

Following digestion, scRNA-seq and flow cytometry analyses demonstrated that this procedure successfully isolates diverse cell types, including epithelial and immune cells. Enzyme solutions should be freshly prepared on the day of the experiment. All digestion parameters are optimized for a single whole murine lung. When using only a single lobe or multiple lungs, the digestion time should be adjusted to ensure high cell viability and complete tissue dissociation. Ambient RNA contamination and multiplet formation are two major factors that affect scRNA-seq quality ^18^. Although dead cell removal kits are often used to enhance viability, this protocol consistently yields over 85% live cells after digestion, making this step optional. The barcode rank plot confirmed minimal ambient RNA contamination even without dead cell removal.

Although this protocol produces high quality single cell suspensions at low cost, process induced cellular stress may still occur during enzymatic digestion. To minimize stress-related gene expression artifacts, a single operator should process no more than four samples simultaneously.

## ACKNOWLEDGMENTS

This work has been supported by National Institute of Health, The National Eye Institute (NEI), USA, grants 5R01EY026202, R01EY035333; the National Institute of Dental and Craniofacial Research (NIDCR) grants R01DE031044, 2R01DE014756, and National Cancer Institute (NCI) grants 5R01CA271500 and 5R01CA211187.

